## Supplemental Figures for "Human birth tissue products as a non-opioid medicine to inhibit post-surgical pain": ELIFE Supplemental Figures 7.2.docx

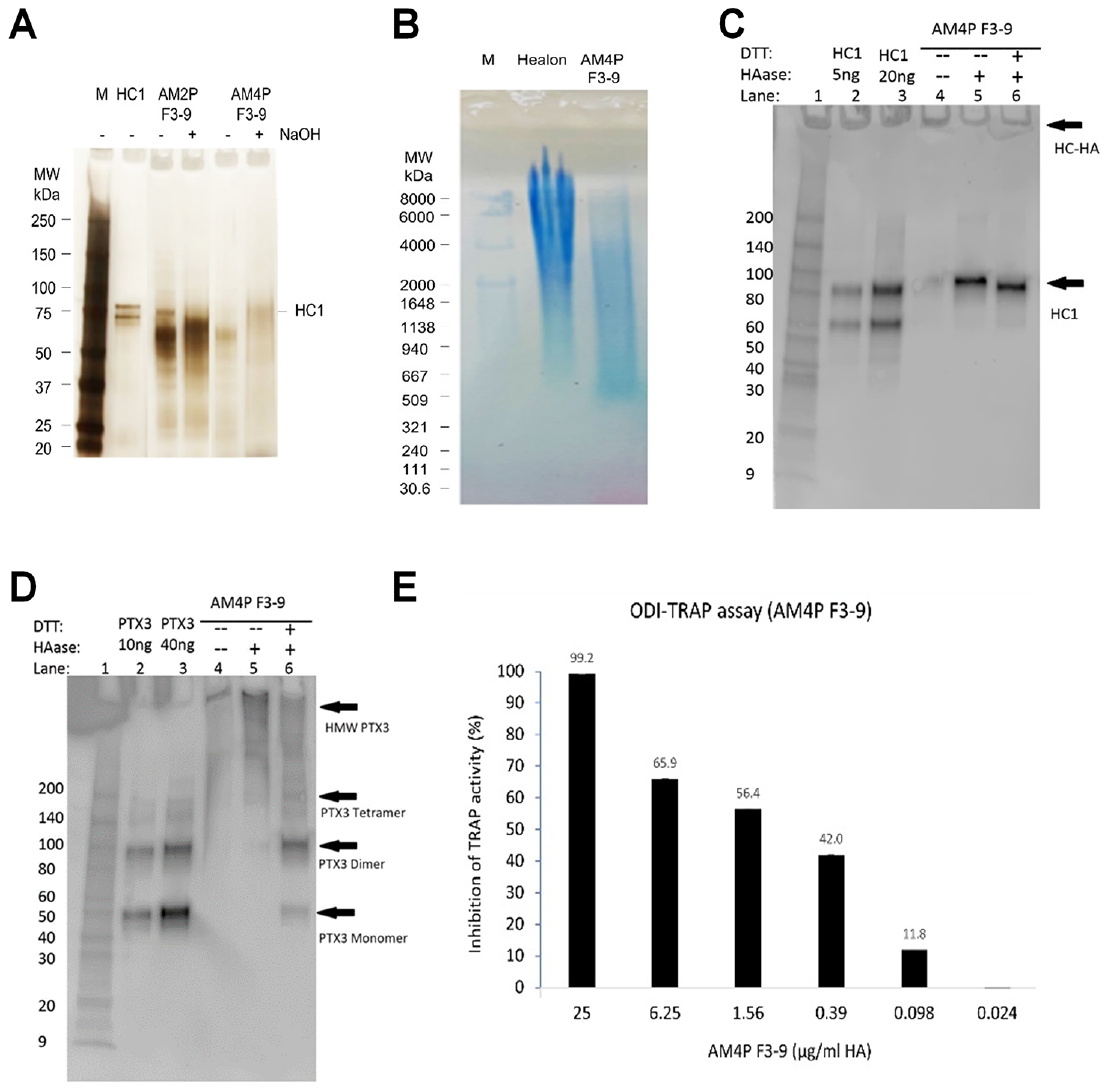

**Fig. S1. Purification and characterization of HC-HA/PTX3.** (A) HC-HA/PTX3 was prepared from the human amniotic membrane by 2 runs (AM2P F3-9) or 4 runs (AM4P F3-9) of CsCl/4 M GnHCl ultracentrifugation and was then analyzed by silver staining. Each lane was loaded with 0.25 µg of HA without or with 100 mM NaOH treatment (25°C, 1 h) to cleave the bond between HA and HC1. (B) HC-HA/PTX3 (AM4P F3-9) purified from the amniotic membrane was electrophoresed on 0.5% agarose gel and stained with All-stains dye. Healon as a high-molecular-weight (HMW) HA control and HC-HA/PTX3 were loaded at 10 µg HA/lane; M: HA molecular weight ladder. (C-D) HC1 and PTX3 in HC-HA/PTX3 were detected by western blot using respective antibodies without (-) or with (+) hyaluronidase (HAase) treatment to release HC1 or HMW-PTX3, of which the latter can then be resolved into dimer or monomer without (-) or with (+) reduction with DTT. (E) Cloned murine RAW264.7 monocytes were seeded at 1x104 cells/cm2 in MEM α/10% FBS and differentiated into multi-nucleated osteoclasts with 25 ng/ml RANKL as the positive control and treated with HC-HA/PTX3 at different concentrations (0.024 – 25 µg/mL) for 3 days. The inhibition of TRAP activity in cell lysates was calculated as a percentage (%) of that of the positive control (shown on the top of each bar).

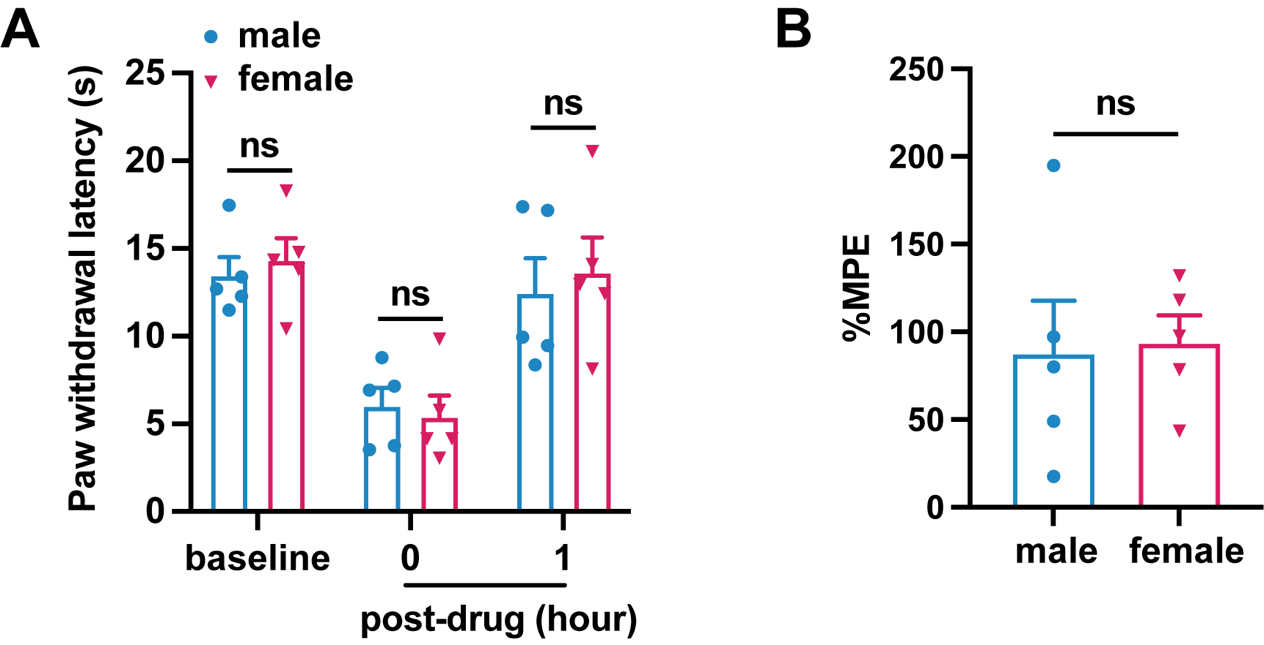

**Fig. S2. HC-HA/PTX3 induced comparable inhibition of heat hyperalgesia in male and female mice after plantar incision.** (A) Paw withdrawal latency at different time point of male and female mice included in Figure 3B. (N= 5/sex) (B) The percentage of maximal possible effects (%MPE) at 1 hour post-drug were calculated for male and female mice included in Figure 3B. %MPE=1 - (baseline - post-drug) / (baseline - pre-drug). Data are mean ± SEM. (A) Two-way mixed model ANOVA followed by Bonferroni post hoc test. ^*^P<0.05 versus male. (B) Unpaired t-test.

**Fig. S3. HC-HA/PTX3 did not affect the excitability of large-diameter DRG neurons in wild-type (WT) mice after the plantar incision.** (A) The representative trace of membrane potential recorded under current-clamp conditions before and after HC-HA/PTX3 (15 µg/mL) treatment in a large DRG neuron of WT mice. Neurons were categorized according to cell body diameter as <20 μm (small), 20–30 μm (medium), and >30 μm (large). (B) The resting membrane potential (RMP) in large DRG neurons was not significantly changed at 5 min after HC-HA/PTX3 (10 μg/mL) treatment, compared to pre-drug (P=0.41). N=5. (C) Representative traces of rheobase measurements before and after HC-HA/PTX3 (10 μg/mL). (D-H) Quantification of the rheobase (D, P=0.09), action potential (AP) threshold (E, P=0.78), AP amplitude (F, P=0.8), AP duration (G, P=0.41), and the mean input resistance (Rinput, H, P=0.17) before and at 5 min after HC-HA/PTX3 (10 μg/mL) treatment. N=5. Data are presented as mean ± SEM. (B, D-H) Paired t-test.

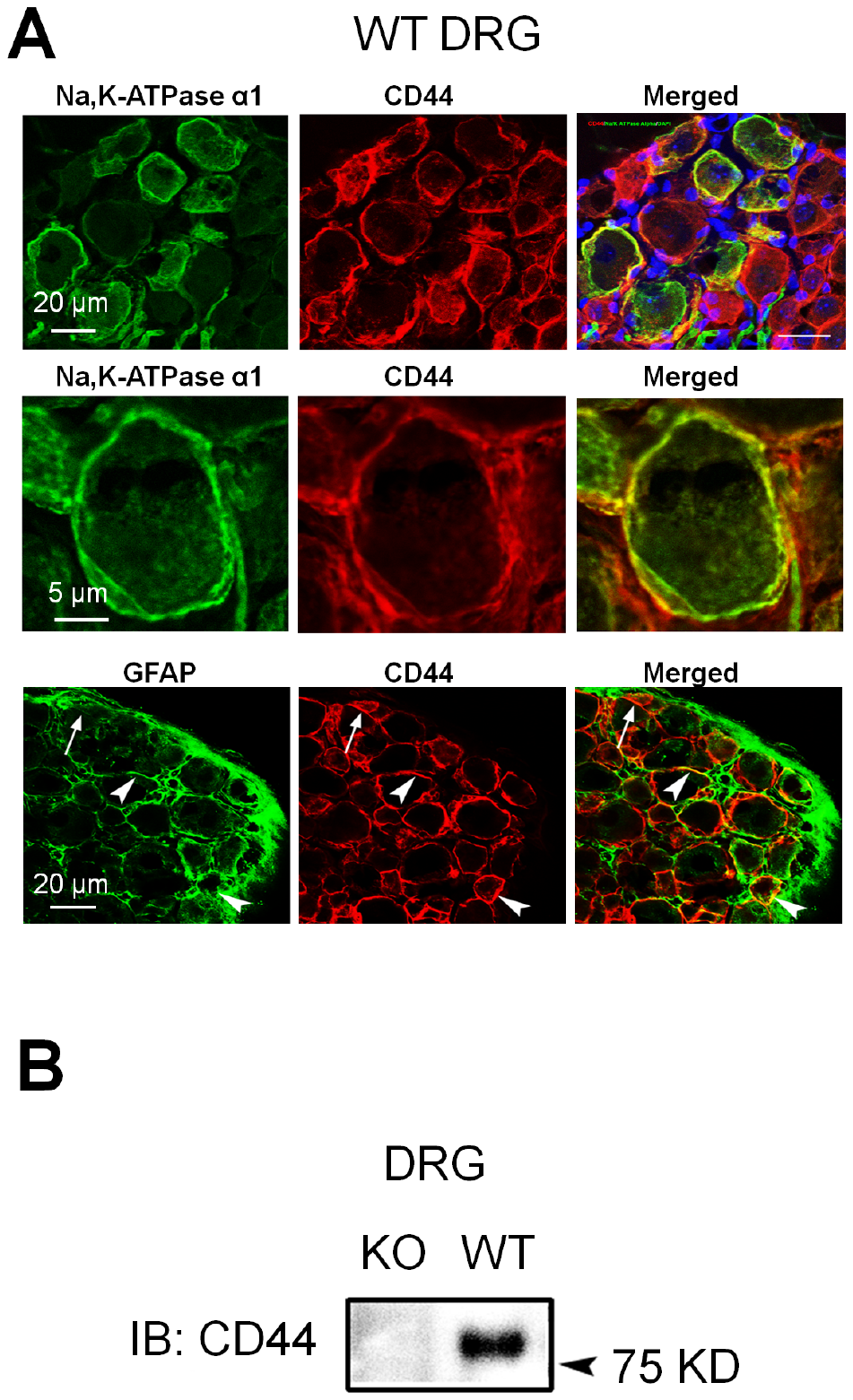

**Fig. S4. The expression of CD44 in mouse DRG.** (A) Upper: Colocalization of CD44 and Na, K-ATPase alpha1 (a neuronal marker) immunoreactivity (IR) in wild-type (WT) mouse DRG. Blue: DAPI. Middle: A higher power view of CD44 and Na, K-ATPase alpha1 colocalization. Lower: Colocalization of CD44 and GFAP (a satellite glial cell marker) in DRG. Arrow: single-labeled cell; Arrowhead: double-labeled cell. (B) The specific CD44 band was not observed in the protein extracts derived from DRG tissues of CD44 knockout (KO) mice in the western blot study.

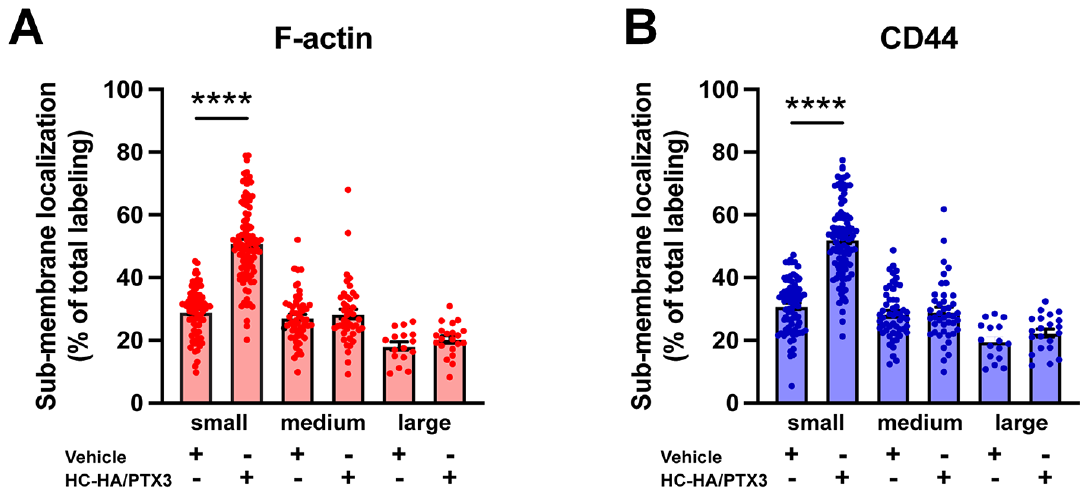

**Fig. S5. Differential effects of HC-HA/PTX3 on sub-membranous F-actin polymerization and translocation of CD44 in small-, medium-, and large-diameter wild-type (WT) DRG neurons.** (A) Quantification of sub-membranous F-actin polymerization and (B) translocation of CD44 in different sizes of WT DRG neurons after HC-HA/PTX3 (10 μg/mL) or the vehicle (saline) treatment. DRG neurons were categorized according to cell body diameter as <20 μm (small), 20–30 μm (medium), and >30 μm (large). HC-HA/PTX3 increased sub-membranous F-actin polymerization and translocation of CD44 exclusively in small neurons. N=16-91/group. Data are mean ± SEM. One-way ANOVA followed by Bonferroni post hoc test. ^****^P< 0.0001 versus vehicle.

**Fig. S6. Quantification of submembranous F-actin polymerization and translocation of CD44 in small-diameter wild-type (WT) DRG neurons in each group.** (A) DRG neurons were electroporated with siRNAs specifically targeting *Ank2* and *Ank3* (siAnk), (B) and those targeting *Ezr*, *Rdx*, and *Msn* (siERM) complex. Neurons were treated with a bath application of vehicle (saline) or HC-HA/PTX3 (10 μg/mL) for 45 min. N=59-114/group. (C) The mRNA expression of *Ank2*, *Ank3*, *Ezr*, *Msn*, and *Rdx* in DRG neurons electroporated with specific siRNAs were assayed by qPCR. N=2. Data are mean ± SEM. One-way ANOVA followed by Bonferroni post hoc test. ^***^P<0.001 versus vehicle; ^###^P<0.001 versus indicated group.

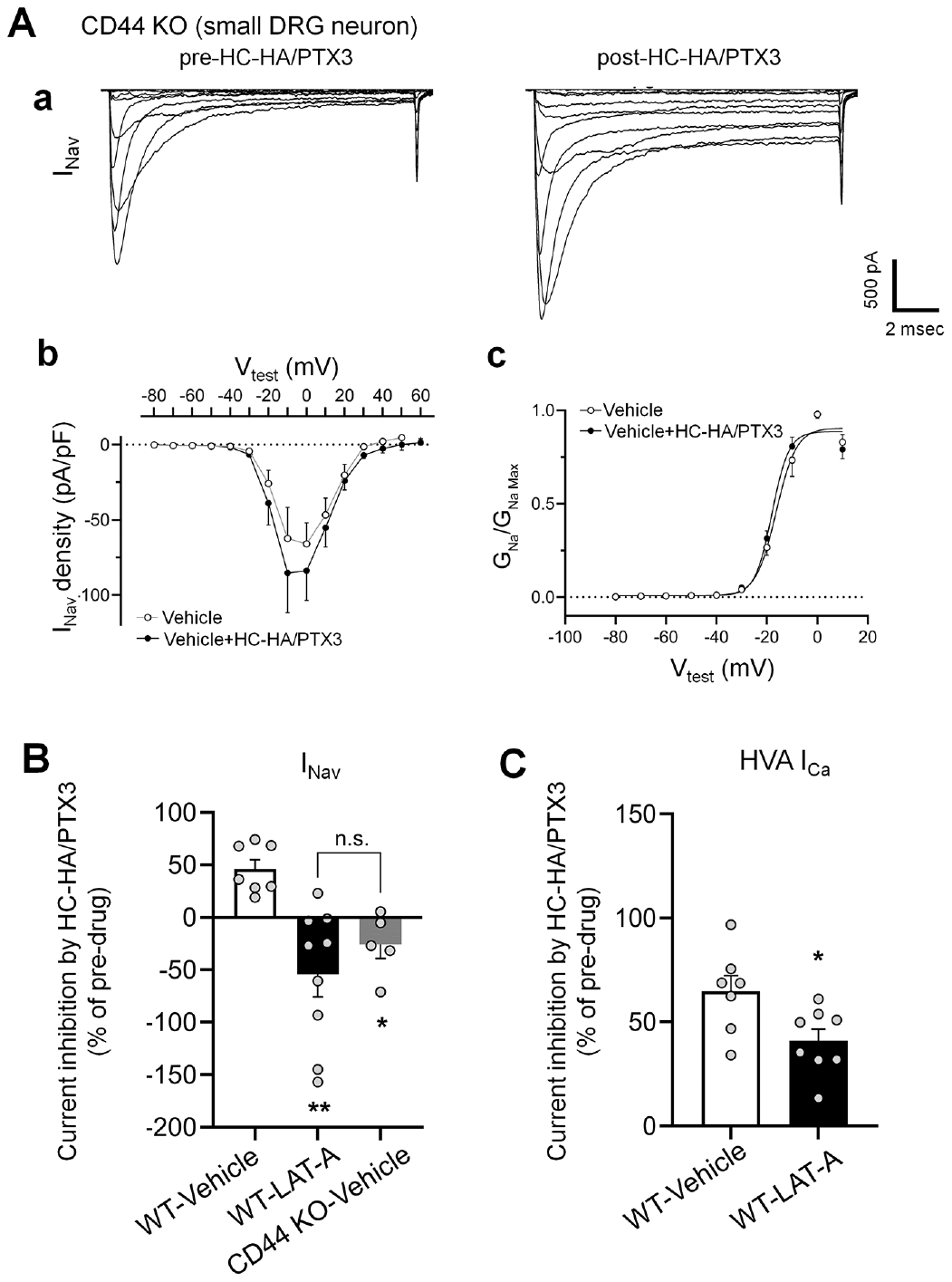

**Fig. S7. The inhibitions of I_Nav_ by HC-HA/PTX3 in small-diameter DRG neurons were diminished in CD44 KO mice and were inhibited by a pre-treatment of LAT-A in neurons from WT mice.** (A) HC-HA/PTX3 (10 µg/mL) did not inhibit I_Nav_ in small-diameter DRG neurons from CD44 KO mice. a. Representative traces of I_Nav_ in a CD44 KO neuron before and at 5 min after bath application of HC-HA/PTX3 (10 µg/mL). b. There was no significant change of I_Nav_ density (pA/pF) before and after HC-HA/PTX3 (10 µg/mL) treatment in CD44 KO neurons. c. HC-HA/PTX3 did not alter G_Na_/G_Na_ max across the test voltages in CD44 KO neurons. N=5. (B) Changes of I_Nav_ after bath application of HC-HA/PTX3 (10 µg/mL) in vehicle-infused (N=7) and LAT-A-infused small WT DRG neurons (P<0.01, N=9), and in vehicle-infused CD44 KO neurons (N=5). DRG neurons were infused with the vehicle or LAT-A (0.5 nM) through the recording electrode, followed by bath application of HC-HA/PTX3 (10 µg/mL) 5 min later. The lumbar DRG neurons were harvested on Days 2-3 after the plantar incision. DRG neurons were categorized according to cell body diameter as <20 μm (small), 20–30 μm (medium), and >30 μm (large). Data are mean ± SEM. One-way ANOVA with Holm-Sidak post-test. ^*^P < 0.05, ^**^P<0.01 versus WT-vehicle pretreatment group. (C) The inhibition of HVA I_Ca_ by HC-HA/PTX3 (10 µg/mL) in vehicle-infused (N=7) and LAT-A-infused small WT DRG neurons (P=0.02, N=8). Data are mean ± SEM. Unpaired Student’s t-test.

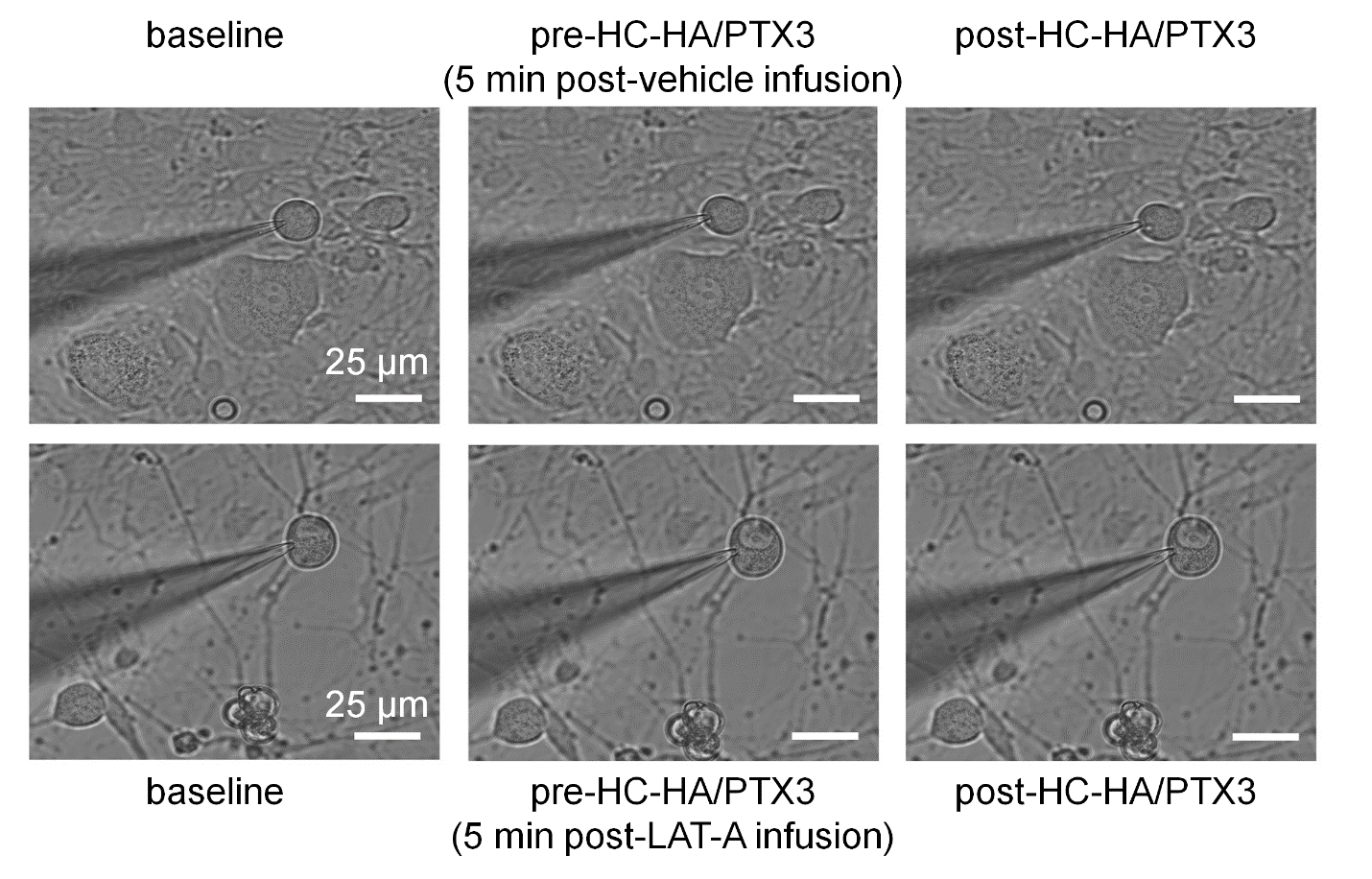

**Fig. S8. Intracellular infusion of LAT-A did not change the gross morphology of DRG neurons in patch clamp recordings.** Example images show a small-diameter DRG neuron after infusion vehicle or Latrunculin A (LAT-A, 0.5 nM) through the recording electrode, followed by bath application of HC-HA/PTX3 (15 μg/mL). Scale bar: 25 µm. DRG neurons were categorized according to cell body diameter as <20 μm (small), 20–30 μm (medium), and >30 μm (large).

**Table S1. The measures of intrinsic membrane properties of small-diameter DRG neurons in WT and CD44 KO mice.** Knocking out of CD44 did not significantly alter the intrinsic membrane property of DRG neurons, as compared to that in WT mice.

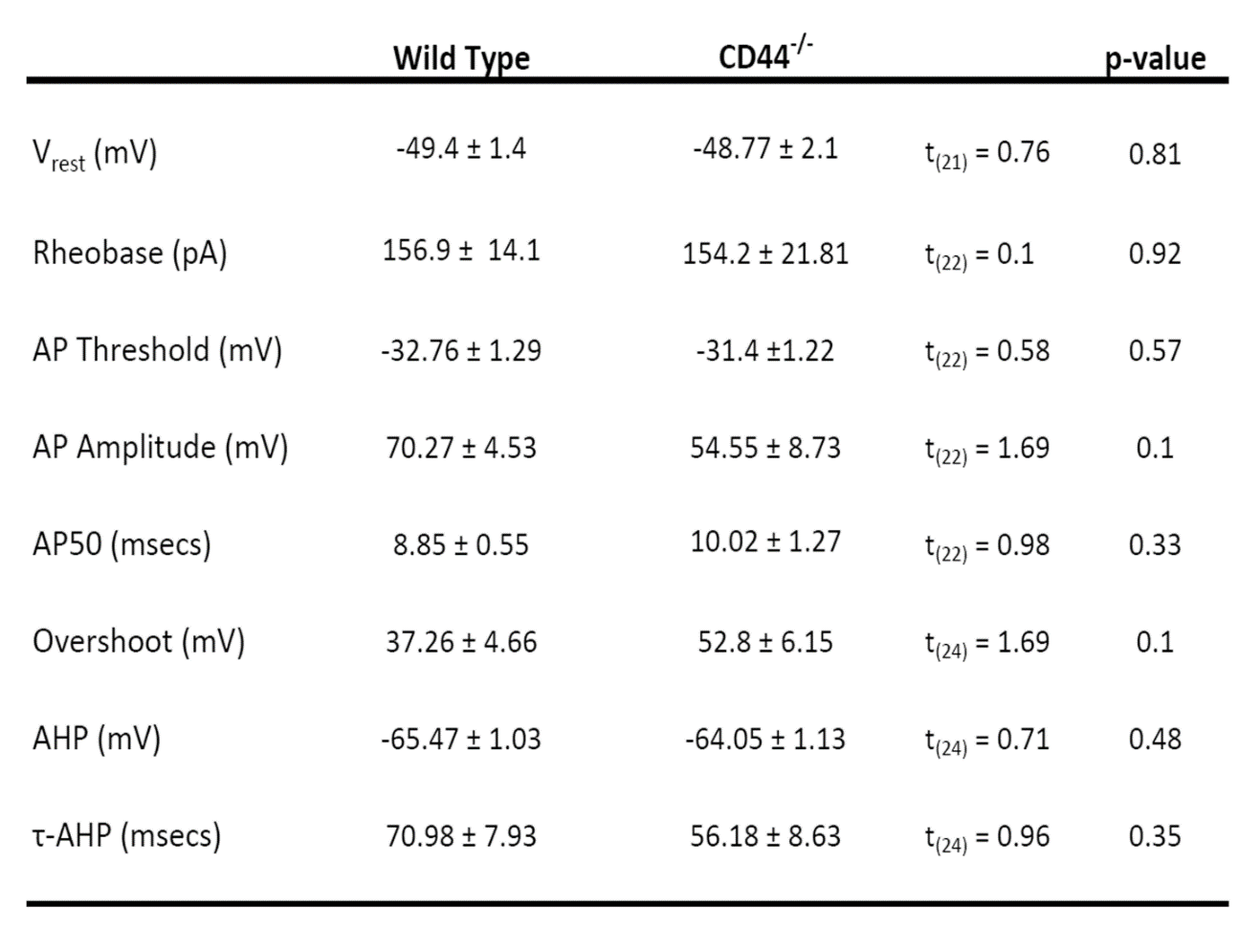
